# Roles of progranulin and FRamides in neural versus non-neural tissues on dietary restriction-related longevity and proteostasis in *C. elegans*

**DOI:** 10.1101/2024.02.06.579250

**Authors:** Dilawar Ahmad Mir, Matthew Cox, Jordan Horrocks, Zhengxin Ma, Aric Rogers

**Affiliations:** Kathryn W. Davis Center for Regenerative Biology and Aging, Mount Desert Island Biological Laboratory, Bar Harbor, ME

**Keywords:** *C. elegans*, Dietary restriction, lifespan, heat shock, proteostasis neurodegeneration, motility

## Abstract

Dietary restriction (DR) mitigates loss of proteostasis associated with aging that underlies neurodegenerative conditions including Alzheimer’s disease and related dementias. Previously, we observed increased translational efficiency of certain FMRFamide-like neuropeptide (*flp*) genes and the neuroprotective growth factor progranulin gene *prgn-1* under dietary restriction in *C. elegans*. Here, we tested the effects of *flp-5*, *flp-14*, *flp-15* and *pgrn-1* on lifespan and proteostasis under both standard and dietary restriction conditions. We also tested and distinguished function based on their expression in either neuronal or non-neuronal tissue. Lowering the expression of *pgrn-1* and *flp* genes selectively in neural tissue showed no difference in survival under normal feeding conditions nor under DR in two out of three experiments performed. Reduced expression of *flp-14* in non-neuronal tissue showed decreased lifespan that was not specific to DR. With respect to proteostasis, a genetic model of DR from mutation of the *eat-2* gene that showed increased thermotolerance compared to fully fed wild type animals demonstrated no change in thermotolerance in response to knockdown of *pgrn-1* or *flp* genes. Finally, we tested effects on motility in a neural-specific model of proteotoxicity and found that neuronal knockdown of *pgrn-1* and *flp* genes improved motility in early life regardless of diet. However, knocking these genes down in non-neuronal tissue had variable results. RNAi targeting *flp-14* increased motility by day seven of adulthood regardless of diet. Interestingly, non-neuronal RNAi of *pgrn-1* decreased motility under standard feeding conditions while DR increased motility for this gene knockdown by day seven (early mid-life). Results show that *pgrn-1*, *flp-5*, *flp-14*, and *flp-15* do not have major roles in diet-related changes in longevity or whole-body proteostasis. However, reduced expression of these genes in neurons increases motility early in life in a neural-specific model of proteotoxicity, whereas knockdown of non-neuronal expression mostly increases motility in mid-life under the same conditions.

## Introduction

Proteostasis is the maintenance of protein synthesis, folding and turnover in the cell and is critical for the normal functioning of tissues. Age-related neurodegeneration is driven or exacerbated by loss of proteostasis, which is frequently characterized by accumulation of toxic unfolded or misfolded proteins^1,2^. Understanding the mechanisms underpinning proteostasis and its disruption in neurodegenerative diseases may help in the development of new therapeutic strategies.

Dietary restriction (DR) is an intervention that mitigates loss of proteostasis associated with aging (see^3^). Benefits of DR, including improved protein processing, require proper adaptation mediated by changes in nutrient sensing. Reduced nutrient availability is transduced through the nutrient-sensing mechanistic target of rapamycin (mTOR)^4^. As a result, signaling to downstream translation factors is reduced, lowering the energetically and materially expensive process of mRNA translation^5^. The redirection of translation to genes that control somatic maintenance is crucial for the health-protective benefits of DR^6^. Thus, while overall translation is lower, translation of mRNAs that preserve cell function is increased relative to those that promote growth ^7–9^. Since many of the health-related phenotypes associated with DR are similar to, or driven by, inhibition of mTOR or translation^4,9,10^, adaptation through differential translation is a key aspect of healthful adaptation to DR conditions.

Restricted access to nutrients, as happens under DR conditions, can lead to changes in behavior and physiology via neuropeptides, which are small signaling proteins that work over a range of temporal and spatial scales^11^. In response to low nutrient availability associated with DR, animals are instinctively driven into a foraging state characterized by increased motility^12^. Previously, we observed changes in total and polysomal (i.e., translated) mRNA via RNAseq from DR involving food dilution and identified genes encoding several neuropeptides that exhibit increased translation efficiency under this condition in *C. elegans*^13^, including Phe-Met-Arg-Phe-NH2 (FMRF)-like peptides, or FRamides.

FRamides genes encode one or more peptides with diversity in their N-terminus that allows them to regulate different biological functions, including neuronal control of stress responses and metabolic regulation^14^. Structurally similar peptides exist from nematodes to more complex systems^15^. In mammals, several groups of FRamides have been identified, with roles in energy metabolism and responding to stress^14,16^. In *C. elegans*, FMRFamide-like peptides (FLPs) comprise the largest and most diverse family of neuropeptides^17^. Expression of *flp* genes is enriched in motor, sensory, and interneurons but is also present in non-neuronal somatic tissues^18,19^.We sought to better understand the roles of some of the more translationally upregulated neuropeptide-encoding *flp* genes under DR^13^, including *flp-5*, *flp-14* and *flp-15*. Although functional information for these neuropeptide-encoding genes is limited, previous researchers showed that *flp-5* and *flp-14* have a stimulatory effect on pharyngeal pumping involved in feeding^20,21^. However, it is unknown whether consequences of altering their expression are limited to behavioral changes or whether they also apply to increased lifespan and proteostatic maintenance normally associated with DR.

In addition to FRamide genes, another secreted peptide translationally increased under DR is progranulin, a growth factor that regulates a number of processes including inflammation, wound healing, metabolism, and others (reviewed in ^22^). Most quiescent cells express progranulin at low levels; however, it is expressed highly in post-mitotic cells of brain and spinal cord. In 2006, it was shown that a single copy mutation of the human form can result in frontotemporal lobar degeneration^23,24^. Low blood levels of progranulin are a biomarker for fronto-temporal dementia (FTD) risk and its functional roles include protecting neurons from premature death^25,26^ and promoting projections from neuronal cell bodies^24^. Raising levels of PGN is being considered as a therapeutic intervention to FTD, but an understanding of its regulation under health-promoting interventions like DR is lacking.

*C. elegans* possesses a homolog of progranulin encoded by *pgrn-1*. The PGRN-1 protein is less complex than its human counterpart, with three granulin domains compared with seven and a half in the human form^27^. Its expression is similar to that of humans, primarily in neurons and intestinal tissue^28^. Interestingly, while mutants have normal lifespan^28^, they exhibit an increased ability to survive certain forms of protein unfolding stress concurrent with an increased rate of apoptotic cell clearance^29,30^. Neuropeptides like FLPs and peptide hormones like PGRN-1 are important mediators of diverse physiological and behavioral functions. In the current study, we investigated the importance of the *pgrn-1* gene as well as neuropeptide genes *flp-5*, *flp-14*, and *flp-15* for lifespan under DR and the survival of DR-conditioned animals under heat stress. Additionally, we utilized a neurodegeneration model that hinders neuromuscular signaling, leading to motility defects^31^, to investigate the influence of these genes on DR-induced adaptation in the presence of neural proteotoxic stress. We explored the impact of low expression of *prgn-1* and *flp* genes on proteostatic maintenance using food dilution and a genetic model of DR via an *eat-2* mutant. The coordination and motility of *C. elegans* are regulated by numerous neurons that innervate body wall muscle cells; misfolded proteins can cause dysfunction in these neurons^32,33^. Motility serves as an indicator of neuronal proteotoxicity and emerges as a novel, aging-dependent health-span marker potentially linked to longevity. Furthermore, we assessed the motility of worms expressing Pan neuronal Amyloid beta (Aβ_1-42_) by measuring their locomotion on solid media. Nematode automated, computer-based motility assay was used as a metric to evaluate Aβ proteotoxicity in *C. elegans*.

## Material and Methods

### Nematode Culture *C. elegans* Strains and lifespan

The *C. elegans* strains were obtained from the *Caenorhabditis* Genetic Center, funded by the NIH Office of Research Infrastructure Programs (R01AG062575). N2 Bristol wild type, and strains mentioned in Supplemental table –S1 were cultured and maintained at 20°C on Nematode Growth Media (NGM) plates seeded with *Escherichia coli* strain OP50 as a food source. Standard methods^34^ were employed for the cultivation and observation of *C. elegans*. All strains underwent growth for at least three generations under normal, unstressed conditions prior to use in experiments. To obtain age synchronized populations for experiments, gravid adult worms laid eggs for 4 hours on OP50 plates before adults were removed and progeny were allowed to develop into day 1 old adults. Except for thermotolerance assays, all experiments were conducted at 20°C. Adult animals were transferred daily to prevent contamination by new larvae. Survival assessments commenced on the 7th day of adulthood, with each strain represented by a population of 100 or more worms distributed across three NGM plates. Survival rates of the animals were assessed daily by observing their movement subsequent to gently tapping the worm using a platinum wire. GraphPad Prism was used for the calculation of mean lifespans and the execution of statistical analyses, with P-values determined using the log-rank (Mantel-Cox) statistic.

### Dietary restriction assays

DR assays were performed as described by Ching and Hsu 2011^35^. For experiments involving RNAi, developmentally synchronous young adult (day 1) worms were transferred to plates containing bacteria expressing dsRNA for 2 days and then transferred to peptone-free nematode growth media plates spotted either Ad libitum (AL) plates (10^11^ CFU/ml OP50) or DR plates (10^9^ CFU/ml OP50). Nematodes were manually transferred every other day to fresh plates until progeny production ceased. Plates were prepared in advance and stored at 4°C. Carbenicillin (25 mg/ml) was included in both AL and DR plates to inhibit bacterial growth. In this study, we have also utilized the *eat-2* mutant, which serves as a valuable genetic model for exploring the complexities of DR.

### Thermotolerance assay

Synchronized populations of day 1 adult *eat-2* mutant worms were transferred to RNAi plates. Nematodes were transferred every day to fresh RNAi plates for seven days. On the seventh day, nematodes were subjected to a heat stress treatment at 35°C for 4 hours, while an unheated control group was maintained at 20°C. Following the heat stress treatment, the recovery of nematodes was assessed over a period of four days by gently tapping the worms with a platinum wire. Survival was measured every day based on their response to touch provocation. Worms that failed to respond to several taps were considered dead and were removed from the plate. Worms that died due to valval rupture, bagging, or crawling off the plate were censored.

### RNAi experiments

Standard bacterial feeding methods were employed for RNA interference (RNAi) studies^36^. Induction of dsRNA was achieved using 1 mM isopropyl β-D-1-thiogalactopyranoside (IPTG). Bacterial clones expressing double-stranded RNA (dsRNA) included the control (empty vector pL4440) sourced from Addgene, Cambridge, MA, U.S.A. All dsRNA sequences targeting various *C. elegans* genes were obtained from the Ahringer RNAi library (Source BioScience, Nottingham, U.K.). *E. coli* HT115 bacteria expressing dsRNA were cultured overnight in LB supplemented with 100 µg/mL Carbenicillin. NGM plates containing 100 µg/mL Carbenicillin and 1 mM IPTG were seeded with the bacteria and allowed to grow for 48 hours before use. All RNAi clones used in experiments underwent sequence verification before use.

### Motility Assays

Two nematode strains were employed in motility assays. The first was the Alzheimer’s model GRU102 strain, which expresses Aβ_1-42_under the *unc-119* promoter for pan-neuronal expression^37^. GRU102 was crossed with a neuron-specific RNAi strain TU3335 to produce ANR201. Motility assessments were performed on the 4th, 7th, and 10th days of adulthood. One hour prior to video recording, animals were transferred to fresh NGM plates without food. For motility assays, young adults were cultured at 25°C. Locomotion data was collected and analyzed using methodology previously described^38,39^. In brief, individual nematodes were tracked continuously in 45-second videos after gently tapping the plates. To assess both forward and backward speed (absolute values) of the nematodes, the videos were analyzed using Wormlab software version 4.1 (MBF Bioscience). For each strain under investigation, 25 videos were captured for subsequent analysis. Differences in motility were determined using the log-rank test in GraphPad Prism 6.0 software.

## Results

### Under DR, *pgrn-1* and *flp* genes do not act in neuronal tissue to promote longevity

In our study, we identified the progranulin gene *pgrn-1* and several *flp* genes that exhibited increased recruitment to polysomes, based on a previously published dataset detailing the transcriptional and translational effects of DR in *C. elegans* (as illustrated in **Fig-S1**). Since these genes are predominantly expressed in neural tissue, we utilized a tissue-specific transgenic *C. elegans* strain (TU3335) engineered to enhance RNA interference capacity exclusively in neurons, with minimal or no RNAi phenotype penetrance in other tissues^40^. Our objective was to investigate whether reducing the expression of these genes in neurons would impact the ability of animals to extend their lifespan under DR conditions. Surprisingly, downregulating the expression of *pgrn-1* or *flp-5*, -14, or -15 genes under DR conditions doesn’t change lifespan compared to the DR animals on control RNAi. These findings suggest that precise regulation of *prgn-1* and *flp* genes in nervous system might not be necessary for the observed lifespan extension under DR conditions (**Fig-1**).

**Figure 1:**
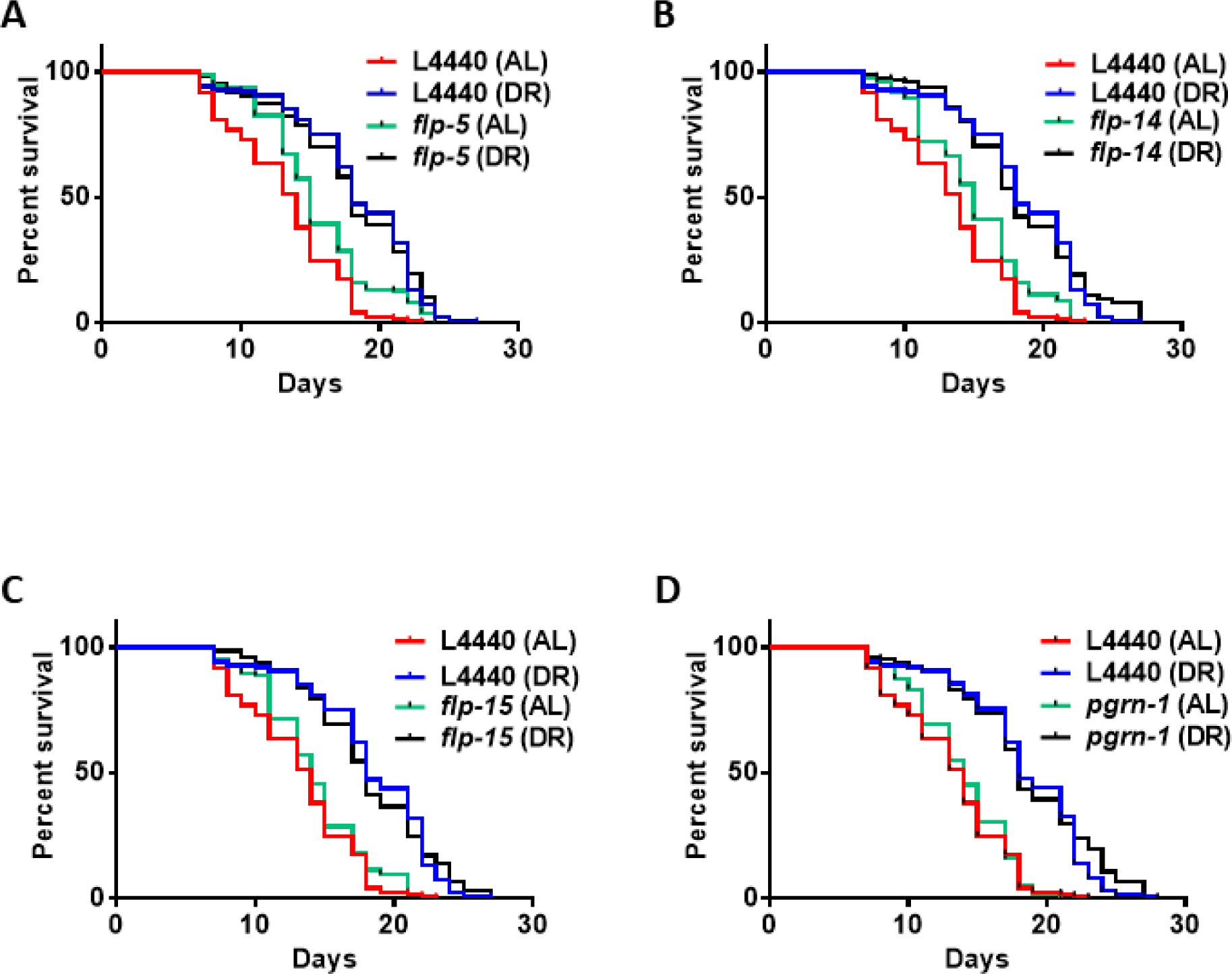
Lifespan of TU3335 treated with *pgrn-1* and *flp* RNAi under AL and DR conditions: TU3335, a neuronal-specific RNAi strain, was fed with control (L4440 empty vector) and bacteria expressing dsRNA specific for *pgrn-1* and FRamides genes (*flp-5*, -14 and -15) for 48 hours starting on the first day of adulthood. On day 3 of adulthood, nematodes were transferred to AL or DR plates for their entire lifespan (**A-D**). (**A**) Under DR conditions, *pgrn-1* knockdown shows no increase in lifespan compared to the control DR (0%, n = 169). The median percentage changes in lifespan from the log-rank test comparing with DR controls are as follows: (**B**) *flp-5* (0%, n= 175), (**C**) *flp-14* (0%, n= 168), (**D**) *flp-15* (0%, n= 141). Kaplan-Meier survival curves were compared using the Mantel-Cox log-rank test. Three biological replicates were conducted for this study. The figure displayed is a representative illustration of the collective findings. The results of the three biological replicates are available in Table S2 of the Supporting Information.

To test the effect of modulating their expression outside the nervous system, we assessed RNAi-mediated suppression targeting both *pgrn-1* and *flp* genes in wild-type N2 animals. Under both DR and AL conditions, we observed a decrease in lifespan specifically associated with RNAi of the *flp-14* gene. However, since DR is still able to increase lifespan when expression of *flp-14* is reduced (P < 0.0001), it does not specifically interfere with the effects of DR. Results indicate that somatic expression of *flp-14* is generally important for normal lifespan in *C. elegans*. Conversely, no statistically significant changes in lifespan were evident in response to RNAi treatment targeting *pgrn-1*, *flp-5*, and *flp-15* (**Fig-2** and details in Table S3 in the Supporting Information). Taken together with results using the neuron-specific RNAi strain TU3335, results indicate that these genes do not play pivotal roles in modulating lifespan extension within the context of DR.

To further investigate whether there is any role for *pgrn-1* and *flp* genes in longevity benefits conferred by DR, we adopted a genetic approach to utilizing a well-established pharyngeal pump mutant, *eat-2(ad465)*, known for its diminished feeding efficiency. RNAi of *flp-14* in the *eat-2* mutant shorten lifespan, like results in Fig. 2. However, there were also significant negative impacts to lifespan of *eat-2* mutants when subjected to RNA interference targeting the *pgrn-1*, *flp-5*, and *flp-15* genes (**Fig-S2** and Table S4 in the Supporting Information for details). It is possible that there is better penetrance of RNAi in *eat-2* mutants than in N2, since they are continuously fed under RNAi conditions (see Methods). This assay highlights the potential importance of these genes in regulating the normal lifespan of nematode worms. It suggests that the proper functioning of these genes may be essential for the nematodes to achieve a full and healthy life under DR conditions.

**Figure 2:**
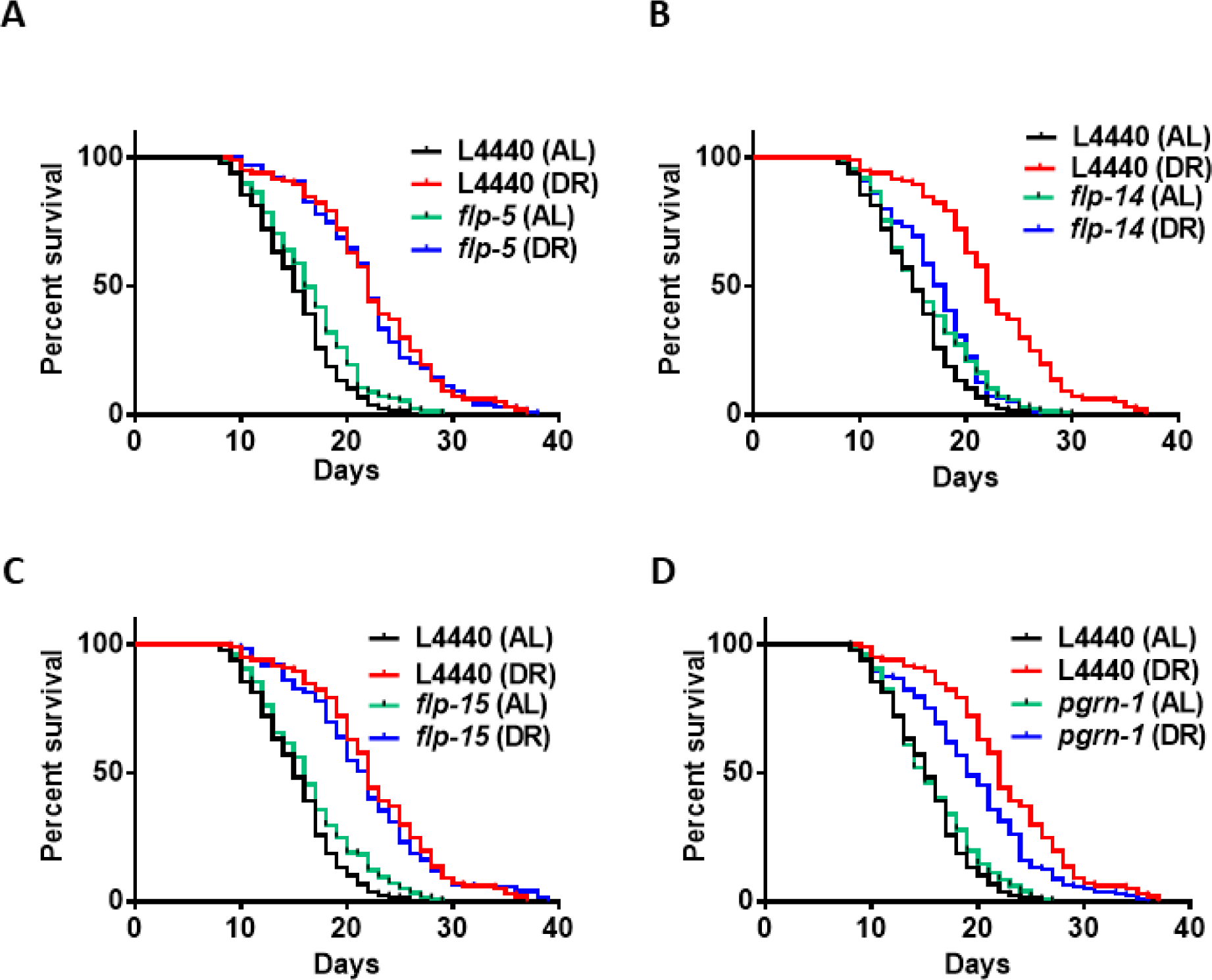
Lifespan of Wild-type treated with *pgrn-1* and *flp* RNAi under AL and DR conditions: Wild-type N2 *C. elegans* were fed dsRNA corresponding to *pgrn-1*, FRamides genes, or the control vector L4440 for 2 days starting from day 1 of adulthood. On day 3 of adulthood, the nematodes were transferred to AL or DR plates for their entire lifespan (**A-D**). The knockdown of *pgrn-1* and FRamides genes has decreased the lifespan of animals compared to the control DR. The median percentage change in lifespan and P-values from the log-rank test comparing with DR controls are as follows: (**A**) *pgrn-1* (-13.63%, n =137), (**B**) *flp-5* (0%, n = 99), (**C**) *flp-14* (-18.18%, n = 111), (**D**) *flp-15* (0%, n = 122). Kaplan-Meier survival curves were compared using the Mantel-Cox log-rank test. The results of the two biological replicates are available in Table S3 of the Supporting Information.

### *pgrn-1* and *flp* genes are not required for maintaining proteostasis in *eat-2* mutants

Elevated ambient temperatures can disrupt the normal structure and function of biological molecules, significantly impacting the overall physiology of animals. In the case of *C. elegans*, elevated temperatures can induce various cellular abnormalities, including neural degeneration and heat-induced necrotic cell death ^,41,42^. In *C. elegans*, from the initial days of adulthood there is a rapid decline in the capacity to uphold proteostasis^43,44^. The primary objective of this assay was to investigate the involvement of the *pgrn-1* and *flp* genes in helping maintain proteostasis in *eat-2* mutant nematodes during the heat shock response (HS).

To elucidate this role, we subjected *eat-2* mutants to 4 hours of heat shock treatment at 35°C, followed by a 24-hour recovery period at 20°C. As expected, the survival rate of the heat-shocked *eat-2* mutants was better than that of control N2 wild type animals (**Fig. 3**). Conversely downregulation of *pgrn-1* and *flp* gene expression had no effect on thermotolerance in the *eat-2* mutant (**Fig-3** and Tables S5 in the Supporting Information). These outcomes suggest that the downregulation of *pgrn-1* and *flp* genes in *C. elegans* are not essential for maintaining proteostasis perturbed by heat stress during DR conditions.

**Figure 3:**
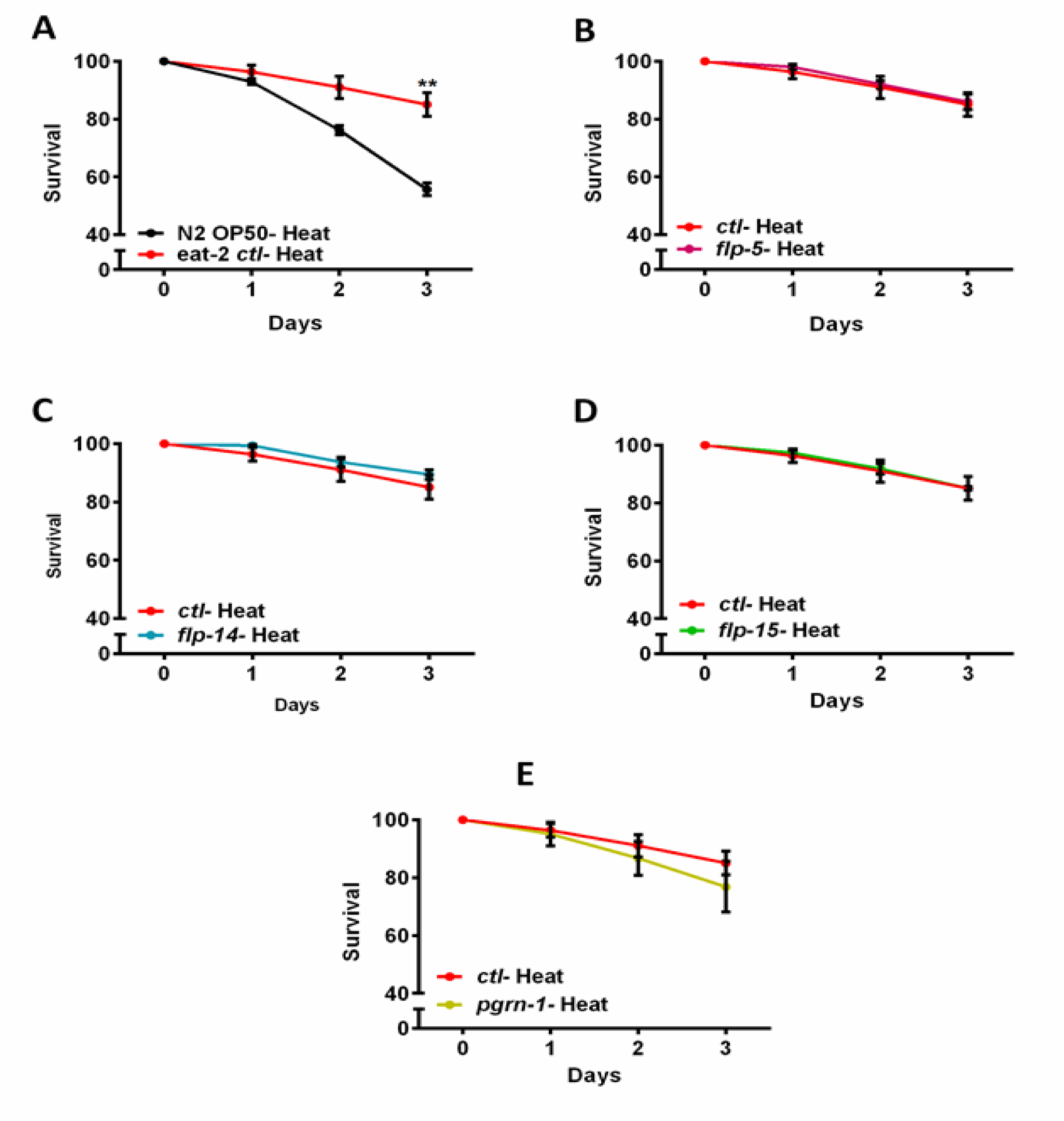
Heat recovery of *eat-2* mutant treated with *pgrn-1* and *flp* RNAi: Bacteria expressing control, *pgrn-1* or FRamides genes dsRNA were fed to animals beginning at day 0 to day 7 of adulthood in **A-E**. Day 7 adult animals were exposed to control conditions (growth at ambient temperature) or thermal stress (35°C for 4 h) on NGM agar plates. Heat-shocked animals were allowed to recover for 12h before scoring for survival. The graph displays the average mean survival up to 3 days’ post heat stress. (**A**) The black line represents the mean resistance to heat stress of wild-type animals (N2). The red lines represent the *eat-2* mutant heat stress in each graph. The *eat-2* mutant exhibited an 85.10% survival after 3 days (n = 121). Compared to control heat-shocked *eat-2* mutant, the percentage heat resistance of *pgrn-1* and *flp genes* RNAi does not show enhanced thermotolerance (**B & E**). The results of the three biological replicates are available in Table S5 of the Supporting Information.

### *pgrn-1* and *flp* genes influence locomotion in an “Alzheimer’s” Model of neurotoxicity

The misfolding and accumulation of proteins have been identified as key hallmarks in various late-onset neurodegenerative conditions such as Alzheimer’s, Huntington’s, and Parkinson’s disease (reviewed in ^45^). The pathogenesis of these neurodegenerative diseases is closely associated with the disruption of protein homeostasis, where the balance between protein production, folding, and degradation is disturbed (reviewed in^44^).

This imbalance results in the accumulation of misfolded or aggregated proteins within neurons, which subsequently leads to cellular toxicity and impaired neural function (reviewed in ^46,47^). Nematode motility is regulated by acetylcholine receptors, which play a crucial role in synaptic functions^48^. The ability to move relies on the maintenance of muscle mass, connective tissue integrity, and proper neuronal signaling^49–53^. Consequently, locomotion serves as a valuable indicator for evaluating neurotransmission. The motility assays conducted on *C. elegans* in our study were devised to investigate the intricate connections between dietary elements, genetic factors, and locomotor behavior. We utilized the Alzheimer’s disease model GRU102 strain, which expresses the Pan-neuronal amyloid beta (1-42) peptide. It is worth noting that GRU102 exhibits impaired neuromuscular and sensorimotor behavior^54^. Our study involved the examination of changes in locomotion at three distinct time points: day-4, day-7, and day-10 of adulthood. We conducted these assessments in both GRU102 and GRU102 crossed with neuronal RNAi strain TU3335 (ANR201) under different dietary conditions. We employed MATLAB worm tracking software to analyze the quantitative data from the video recordings of motility (see Methods). In the GRU102 strain, we observed that knockdown of *flp-14* resulted in decreased motility under AL conditions by day 4 (221.7 vs 169.6 µm/s, P < 0.0001), which then reversed to an increased level compared to both AL and DR control by day 7(115.7 vs 157.7 & 129.5 vs 157.6 µm/s, P < 0.0001). Also, by day 7, *flp-15* RNAi decreased motility under AL conditions (115.7 vs 99.59 µm/s, P < 0.01). Interestingly, *pgrn-1* RNAi decreased motility under AL conditions but increased motility under DR compared with control RNAi under these conditions (115.7 vs 87.69 µm/s, P < 0.0001).

By day 10 of adulthood, both AL and DR animals displayed a dramatic decrease in motility without significant differences between test and control populations. (**Fig-4A**). In contrast, neuronal knockdown of *pgrn-1* and *flp* genes in ANR 201 increased motility by day 4 under AL and DR conditions but had little or no effect at later time points (**Fig. 4B**). These results suggest that downregulating *pgrn-1* and *flp* genes in *C. elegans* are vital for maintaining motility during early stages of life. This downregulation appears to have minimal or no effect on motility during adulthood under DR conditions.

**Figure 4:**
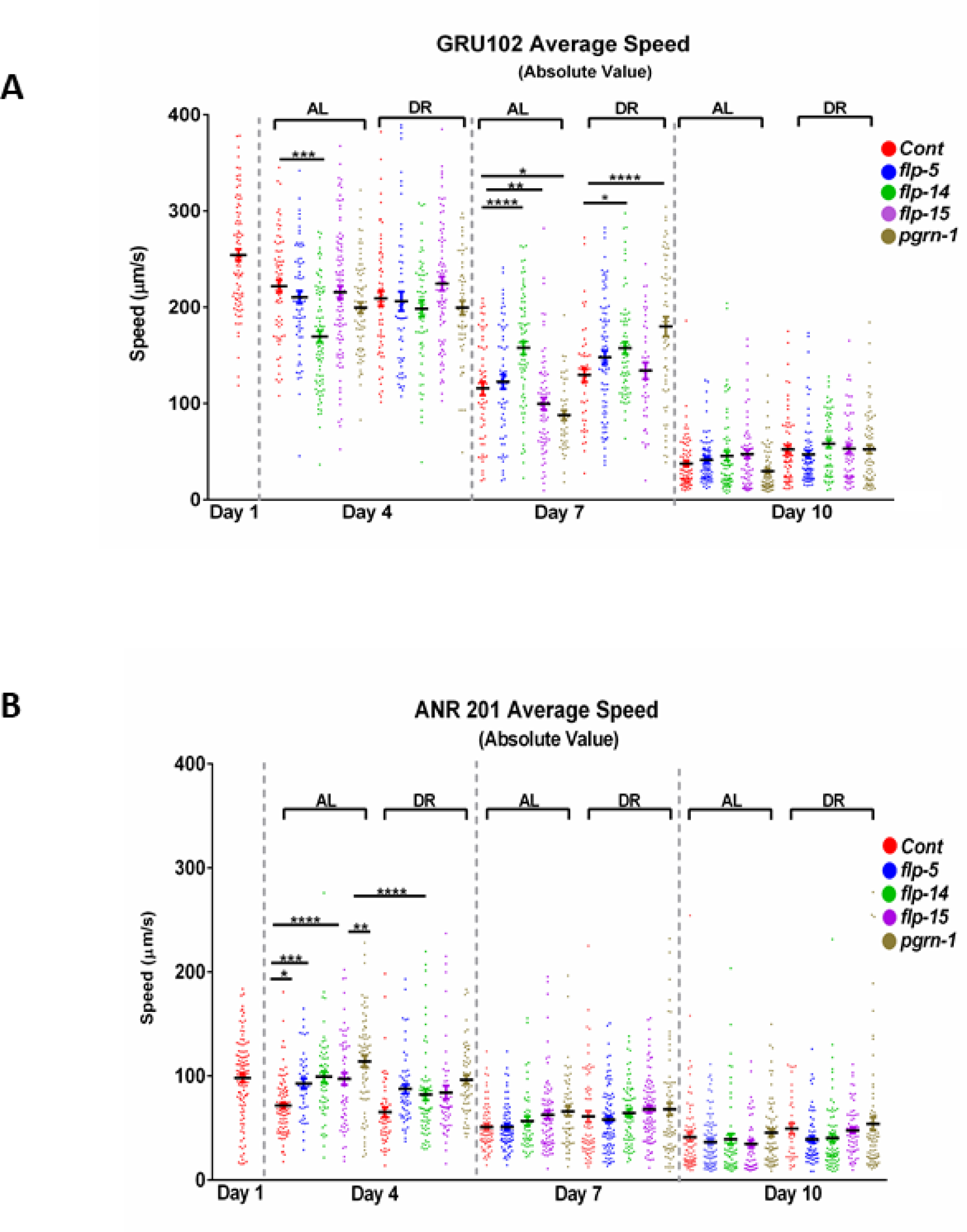
Motility of GRU102 and ANR201 strains treated with *pgrn-1* and *flp* RNAi: Average speed was measured in animals fed bacteria expressing dsRNA for 24 hours starting from day 0 of adulthood for the specified genes. On day 3 of adulthood, the nematodes were placed on AL or DR plates, and the animals were transferred to fresh DR plates for video recording after acclimatization. For both strains, the speed was measured at three different time points (4th, 7th, and 10th day) of adulthood. (**A**) Motility of GRU102 strain expressing Pan-neuronal Aβ_(1-42)_. (**B**) Motility of ANR201 strain. In each experiment, more than ∼70 worms were tracked for 45 seconds. The scatter plots display bars representing the mean and standard error of the mean for all conditions. Statistical comparisons were performed using one-way ANOVA with Dunnett’s multiple comparisons test (*p < 0.05, **p < 0.01, ***p < 0.001, ****p < 0.0001).

## Discussion

This study investigated the role of neuronal and non –neuronal tissues in mediating effects of low *pgrn-1* and *flp* gene expression during normal feeding or DR conditions on *C. elegans* longevity and proteostasis. The latter was tested by thermotolerance and maintenance of motility during adulthood in a neural proteotoxicity model. Results indicate that neither neural expression nor systemic expression of *pgrn-1*, *flp-5*, *flp-14*, and *flp-15* is important for DR-mediated changes in lifespan or thermotolerance. The *flp-14*, while important for normal lifespan, did not prevent DR from extending lifespan. Interestingly, systemic expression of *flp-15* and *pgrn-1* were important for motility under normal feeding conditions in a model of neural proteotoxicity, while knockdown of *pgrn-1* led to increased motility under DR conditions. As opposed to its effect on lifespan, systemic knockdown of *flp-14* increased motility under both AL and DR conditions by day seven of adulthood. Finally, neuronal knockdown of *pgrn-1* and *flp* genes increased early life motility but had little effect in later life.

Previous studies generally support findings of this investigation. the studies showed that, in *C. elegans,* the loss-of-function mutant of *pgrn-1* exhibits a normal lifespan and grossly appears normal^29^. Similarly, overexpression of progranulin does not produce a significant alteration in lifespan^55^. However, during development, the *pgrn-1* functions as a neurotrophic factor and is crucial for neuronal development, regulating neurite outgrowth and enhancing neuronal survival^43^.

The regulation of proteostasis, encompassing the critical processes of protein synthesis, folding, and turnover, is integral to sustaining cellular function and averting age-related neurodegenerative conditions typified by protein misfolding and aggregation. Disturbances in proteostasis have long been associated with the onset of various diseases characterized by protein aggregation. The catastrophic breakdown of cellular proteostasis is implicated as an initial event in the aging process reviewed in ^56^. Therefore, interventions capable of postponing this breakdown demonstrate positive effects on both health and longevity. In our study, we demonstrated the efficacy of DR in delaying proteostasis collapse during adulthood in the *eat-2* mutant compared to the wild type. However, our study indicates that knockdown of *pgrn-1* and *flp* genes don’t play a crucial role in proteostasis associated with thermotolerance. However, their role in maintaining motility in neural proteotoxicity model is more complicated (Fig 3).

In *C. elegans*, the maximum movement speed has been proposed as a highly indicative metric for assessing the health. It shows a strong correlation with longevity and demonstrates the potential to predict lifespan^43^. Additionally, the movement speed of *C. elegans* bears resemblance to the gait speed in humans, establishing it as an essential parameter for understanding health and aging mechanisms in these organisms^57^. Worm-tracking technology used in this study provides detailed access to *C. elegans* motility parameters such as speed, distance traveled, omega bends amplitude in sinusoidal movement, swimming, and thrashing. This study aimed to elucidate locomotion-related behaviors within the GRU102 and ANR 201 strains according to reduced expression of *pgrn-1* and *flp* genes. In a previous study, GRU102 Aβ-expressing young nematodes displayed regular motility, however, their mobility declined from middle age (day 8) onward^33^. We assessed motility in relation to *pgrn-1* and *flp* genes and found that knocking down neuronal expression of these genes increased motility early in life in a manner that does not depend on DR. The most dramatic DR-related effects were observed at day seven of adulthood for *pgrn-1* and *flp-15*. While RNAi of these genes decreased motility under standard feeding, DR reversed effects to increase motility, suggesting that knockdown of these genes made the motility-preserving or enhancing effects of DR even greater than usual.

This study investigated roles of certain neuropeptides (*pgrn-1* and *flp* genes) in extending the healthy lifespan of dietary-restricted *C. elegans*. Future investigations aimed at identifying the cellular and molecular pathways responsible for sensing dietary restriction, regulating distinct aspects of proteostasis, and facilitating communication between neuronal and non-neuronal tissues will be crucial for obtaining a more comprehensive understanding of how neuronal tissues contribute to an organism’s longevity, healthspan, and proteostasis.

## Acknowledgements

This work was supported by the grant from the NIH (R01AG062575-03S1). In addition, research reported in this publication was supported by an Institutional Development Award (IDeA) from the National Institute of General Medical Sciences of the National Institutes of Health under grant numbers P20GM103423 and P20GM104318. Some strains were provided by the CGC, which is funded by NIH Office of Research Infrastructure Programs (P40 OD010440).

## Legend for Supplementary Figures

**Figure S1.**
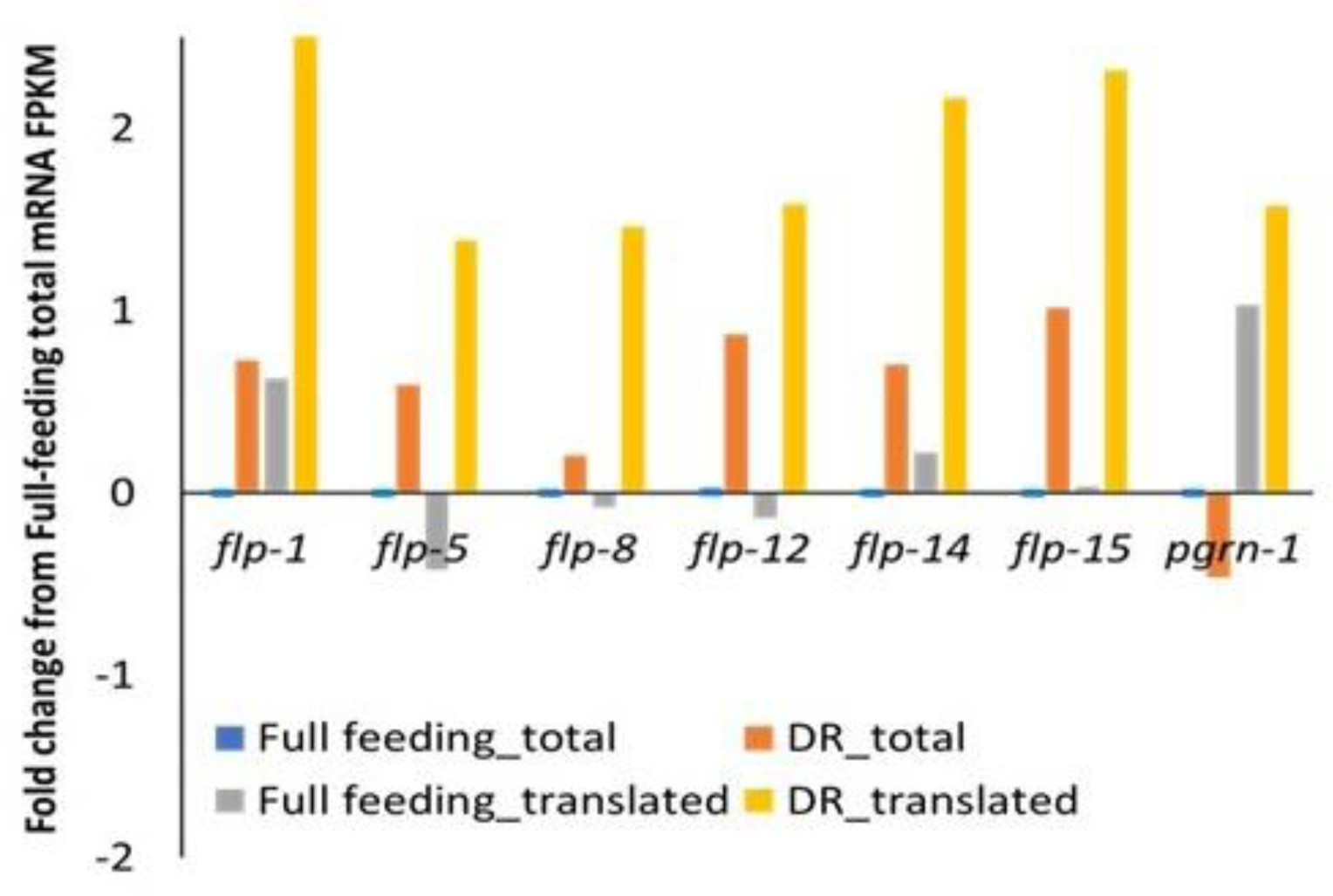
Total versus translated mRNA for progranulin and neurotransmitters under dietary restriction conditions. RNAseq data analysis identifies genes that are translationally regulated under DR. For *pgrn-1* gene under DR results show that translation is upregulated and for *flp* genes both transcription and translation are increased. Fold change in total and translated (polysome-associated) transcript levels compared to full fed control total mRNA levels for the genes shown. All RNAseq experiments were carried out in four biological replicates. Control values used to zero average results. Test is DR animals; control is fully fed. FPKM = Fragments per Kilobase Megabase.

**Figure S2.**
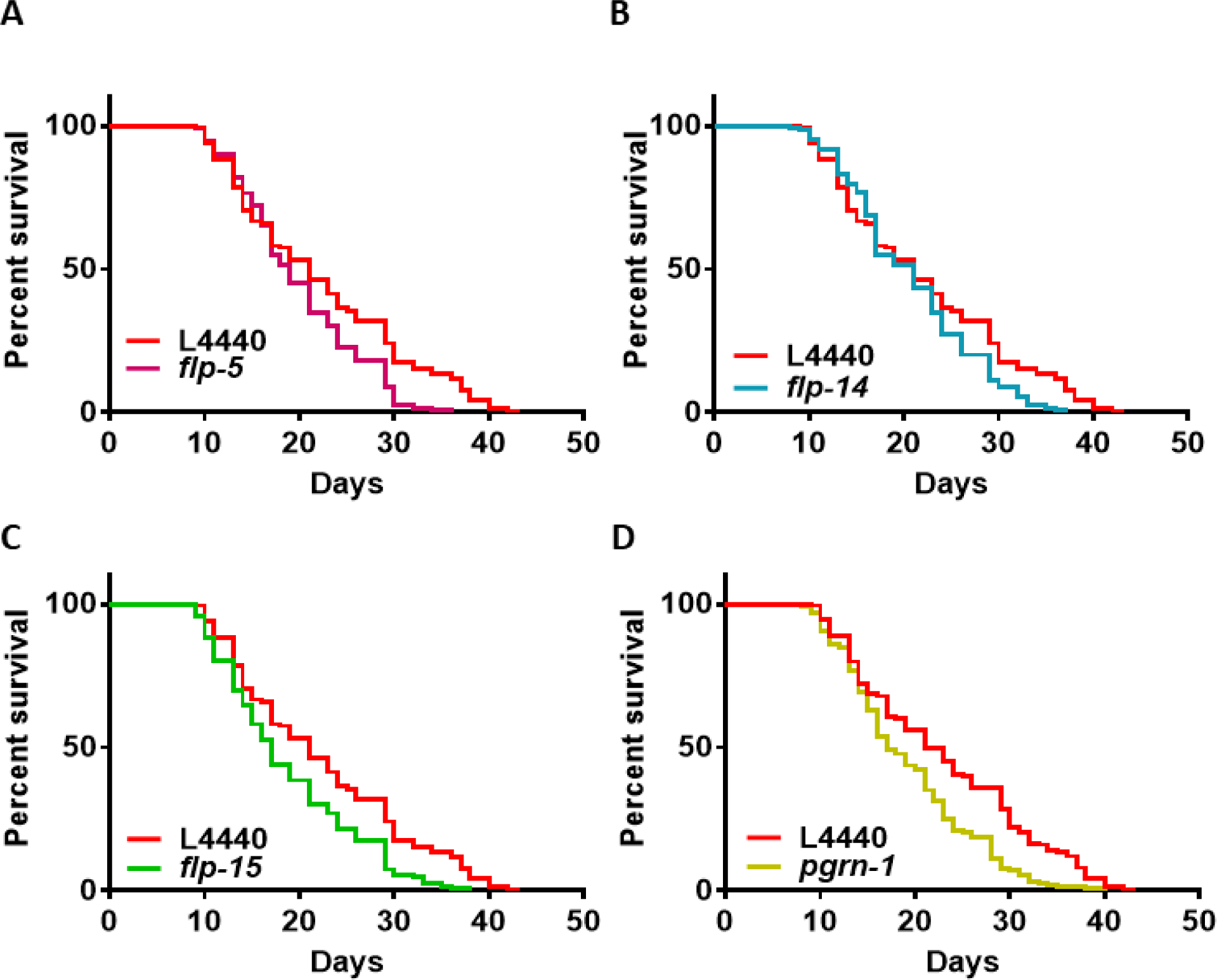
This figure depicts the outcomes of RNAi targeting the *pgrn-1* and *flp* genes in the DA465 strain (*eat-2* mutant) to evaluate their effects on lifespan. The bar graphs demonstrate a detrimental effect on the lifespan of *eat-2* mutants when subjected to RNAi targeting the *pgrn-1* and *flp* genes. Compared to the DR control, knockdown of *pgrn-1* and *flp* genes resulted in decreased lifespan. The percentage survival of minimum lifespan and associated P-values from the log-rank test, comparing with the controls, are as follows (**A**) *pgrn-1*(-19.04%, n = 151), (**B**) *flp-5* (-9.52%, n = 153) (**C**) *flp-14* (0%, n = 124), (**F**) *flp-15* (-19.04%, n = 148). Kaplan-Meier survival curves were compared using the Mantel-Cox log-rank test.

**Figure S3.**
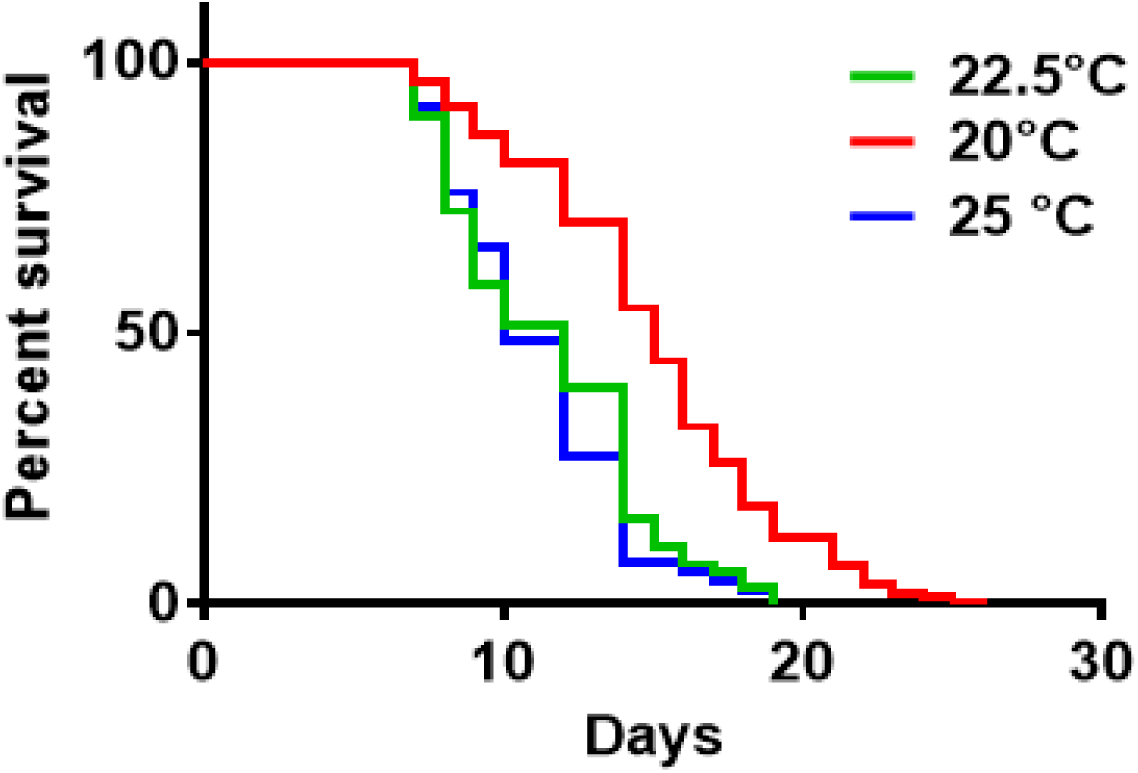
The *C. elegans* strain GRU102 constitutively expresses full-length human Aβ_1–42_, resulting in a notably decreased lifespan and diminished health-span. Kaplan-Meier survival analysis was conducted on Aβ-expressing nematodes at different temperatures, revealing that GRU102 strain nematodes exhibited a significantly reduced lifespan at 25°C compared to those maintained at 22.5°C, while nematodes at 20°C displayed a normal lifespan. This difference was statistically significant (log-rank test: p < 0.001).

**Supplementary Table S1.** List of strains that were used in this study.

| Strain name | Genotype | Backcrossed | Source |
| --- | --- | --- | --- |
| N2 | Wild type | n/a | CGC |
| TU3335 | *lin-15B(n744) X; uls57* | n/a | CGC |
| DA465 | *eat-2(ad465) II* | n/a | CGC |
| GRU102 | *gnals2 [myo-2p::YFP + unc-119p::Abeta1-42]* | n/a | CGC |
| ANR201 | *lin-15B(n744) X; uls57; gnals2 [myo-2p::YFP + unc-119p::Abeta1-42].* | 3x | This study |

**Supplementary Table S2.**
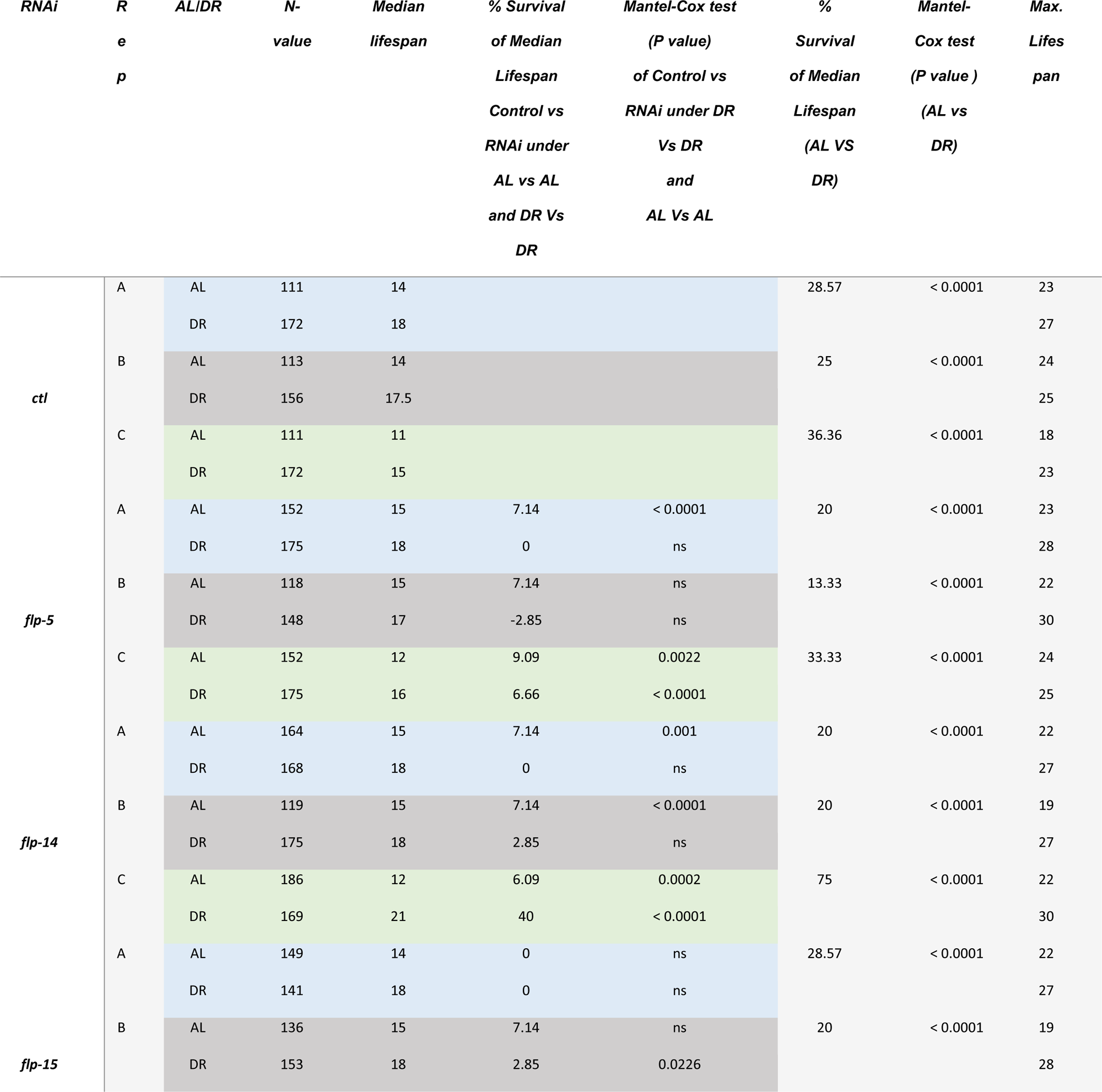
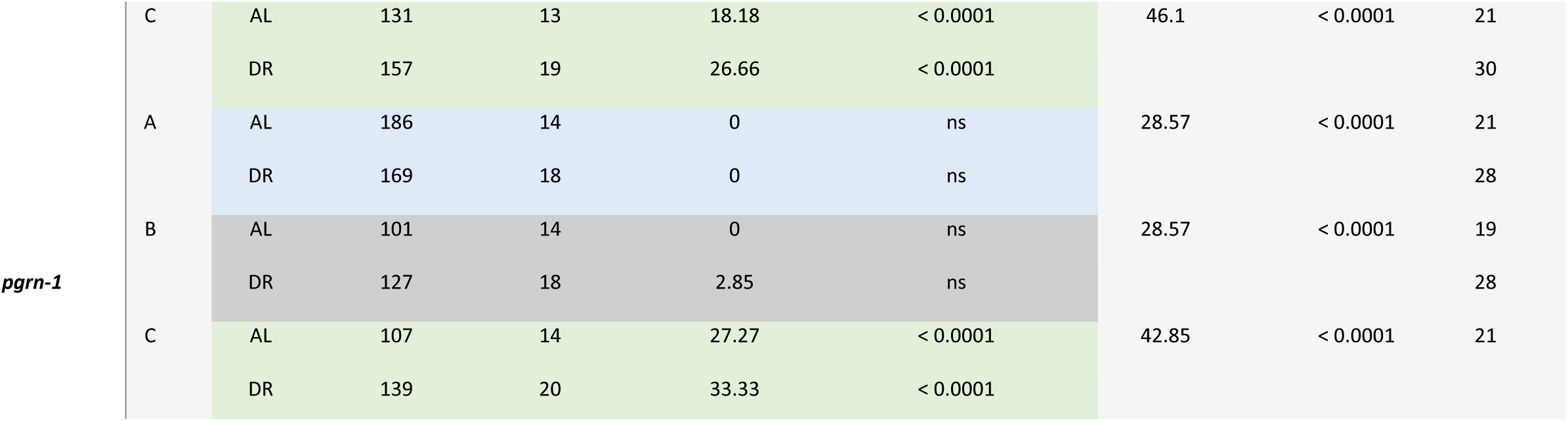
Survival of the TU3335 under AL vs DR.

**Supplementary Table S3.**
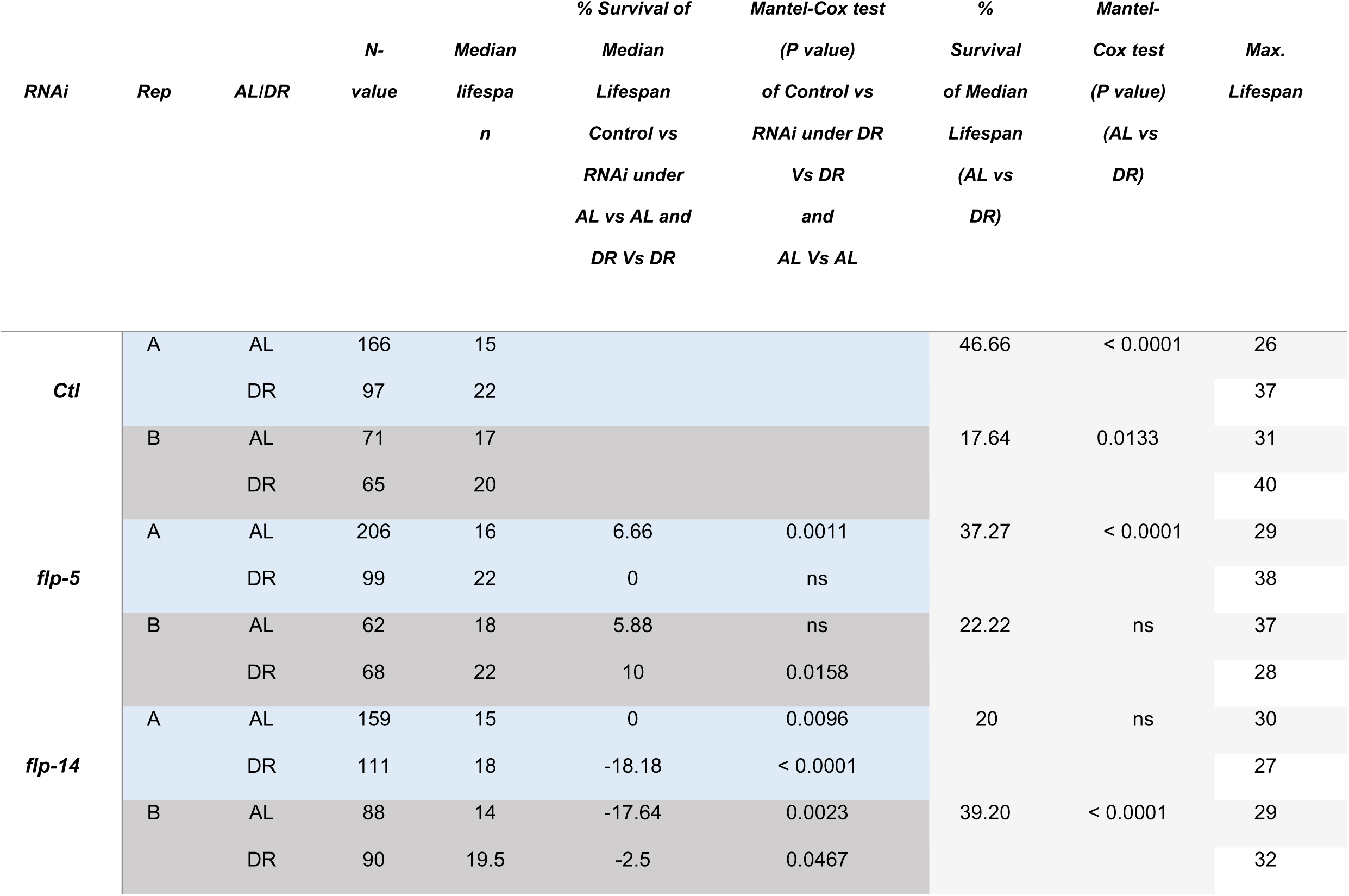
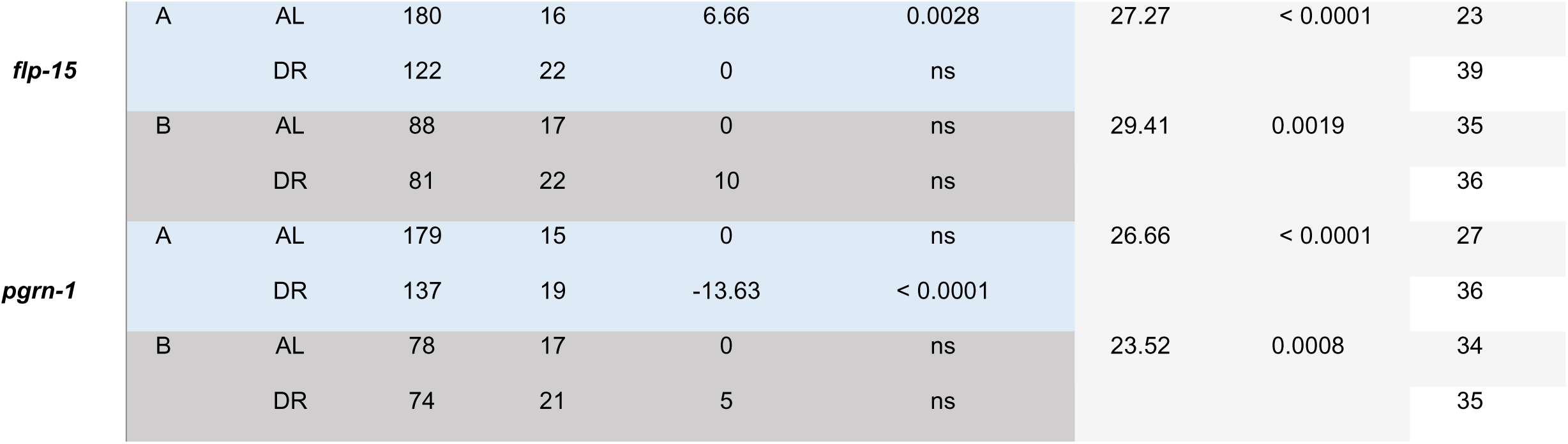
Survival of the N2 Wild type under AL vs DR.

**Supplementary Table S4.** Survival of the *eat-2* Mutant.

| *RNAi* | *N-val* | *Median*<br>*lifespan* | *Max.*<br>*Lifespan* | *% Survival of*<br>*Median*<br>*Lifespan* | *Mantel-Cox*<br>*test (P value)* |
| --- | --- | --- | --- | --- | --- |
| *ctl* | 138 | 21 | 43 |  |  |
| *flp-5* | 153 | 19 | 36 | -9.52 | 0.0003 |
| *flp-14* | 124 | 21 | 37 | 0 | 0.0106 |
| *flp-15* | 148 | 17 | 38 | -19.04 | < 0.0001 |
| *pgrn-1* | 151 | 17 | 40 | -19.04 | < 0.0001 |

**Supplementary Table S5.**
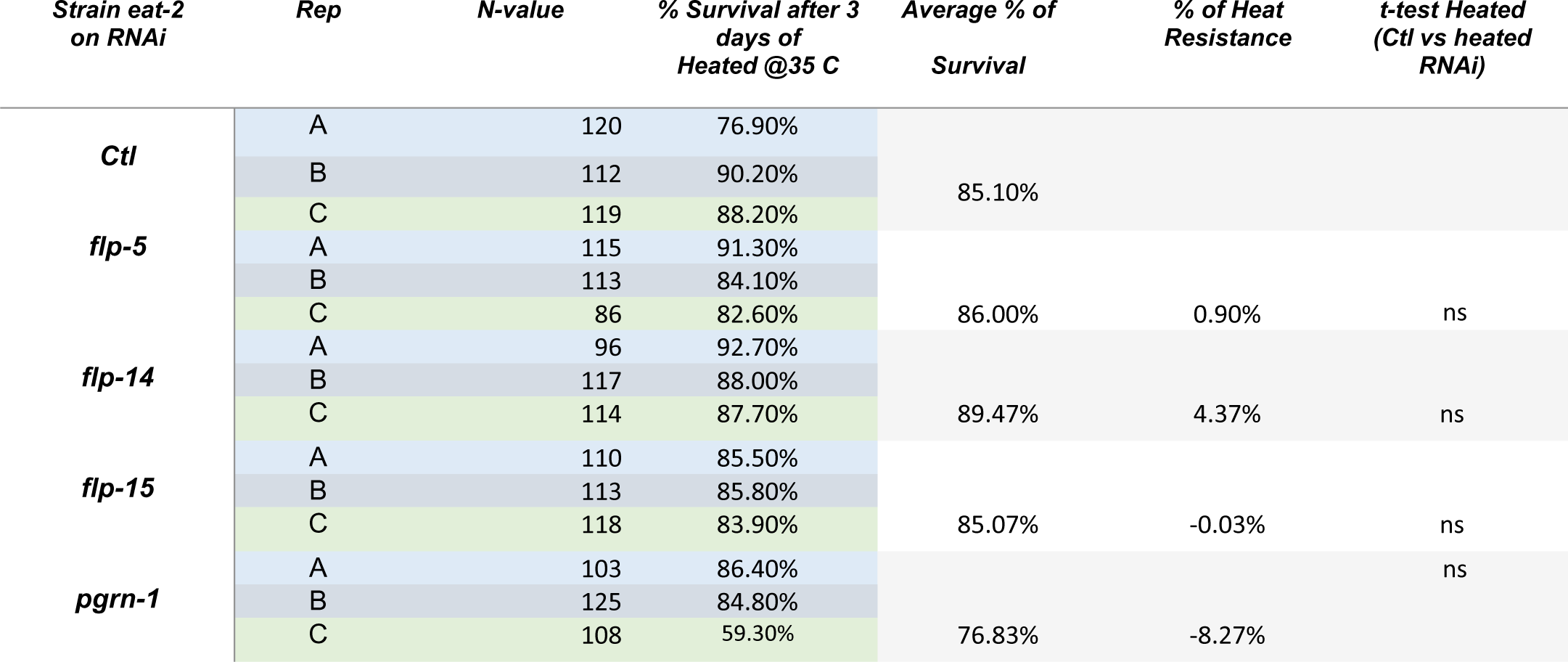
*eat-2* Survival statistics of the heat stress recovery assay.

